# Honest signalling of trustworthiness

**DOI:** 10.1101/2019.12.11.873208

**Authors:** Gilbert Roberts

## Abstract

Trust can transform conflicting interests into cooperation. But how can individuals know when to trust others? Here, I develop the theory that reputation building may signal cooperative intent, or ‘trustworthiness’. I model a simple representation of this theory in which individuals (1) optionally invest in a reputation by performing costly helpful behaviour (‘signalling’); (2) optionally use others’ reputations when choosing a partner; and (3) optionally cooperate with that partner. In evolutionary simulations, high levels of reputation building; of choosing partners based on reputation; and of cooperation within partnerships emerged. Costly helping behaviour evolved into an honest signal of trustworthiness when it was adaptive for cooperators, relative to defectors, to invest in the long-term benefits of a reputation for helping. I show using game theory that this occurs when cooperators gain larger marginal benefits from investing in signalling than do defectors. This happens without the usual costly signalling assumption that individuals are of two ‘types’ which differ in quality. Signalling of trustworthiness may help explain phenomena such as philanthropy, pro-sociality, collective action, punishment, and advertising in humans and may be particularly applicable to courtship in other animals.

## Introduction

When individuals perform costly acts such as helping others, they may in doing so reveal or actively display information about themselves. This information may then induce others to behave preferentially towards them. Building up a good reputation by being seen to be helpful may be adaptive through signalling (*sensu* (Maynard Smith and Harper 2003)); and may have been a major driver of social behaviour especially among humans (Alexander 1987). However, we lack a coherent theory of when investing in a reputation for being trustworthy is adaptive.

One approach to understanding reputation-building is to view it as a strategic investment in making individuals attractive partners (Roberts 1998). This theory, sometimes referred to as reputation-based partner choice (Sylwester and Roberts 2010), postulates that acts such as charitable donations, volunteering, courtship gifts, or contributions to public goods make individuals more attractive partners for relationships. It is based on individuals interacting in social groups and being able to choose among potential interaction partners who differ in partner value by observing their behaviour (Roberts 1998). It is then inferred that those that honestly signal their high quality (Gintis, Smith, and Bowles 2001) will have preferential access through assortative pairing (Johnstone 1997) to mutually beneficial partnerships, which may be social or sexual. This is predicted to lead to an escalation of investment in being seen to be helpful so as to assortatively pair with the most cooperative partners, a phenomenon called competitive altruism (Roberts 1998, Van Vugt et al. 2007, Raihani and Smith 2015, Roberts 2015)

Accumulating evidence from the lab and field supports the role of reputation building in choice of partners (Sylwester and Roberts 2010; Barclay and Willer 2007; Hardy and Van Vugt; Sylwester and Roberts 2013; Raihani and Smith 2015). It is these distinct stages of costly reputation building followed by profiting from partnerships which means that reputation-based partner choice can potentially explain altruism directed unconditionally rather than by direct or indirect reciprocity (Roberts 2015a).

Reputation-based partner choice differs fundamentally from indirect reciprocity as shown in Figure 1 (see also Roberts 2015a). Indirect reciprocity is a process in which one individual incurs a cost and benefits a second individual. A third individual then pays a cost and benefits the original actor, who consequently profits when this benefit exceeds its cost. Indirect reciprocity was discussed by (Alexander 1987), modelled by (Nowak and Sigmund 1998) and demonstrated by (Wedekind and Milinski 2000). However, any form of reciprocity is by definition conditional upon the behaviour of a recipient and so is not considered here as a candidate process underlying the phenomena of interest.

**Figure 1.**
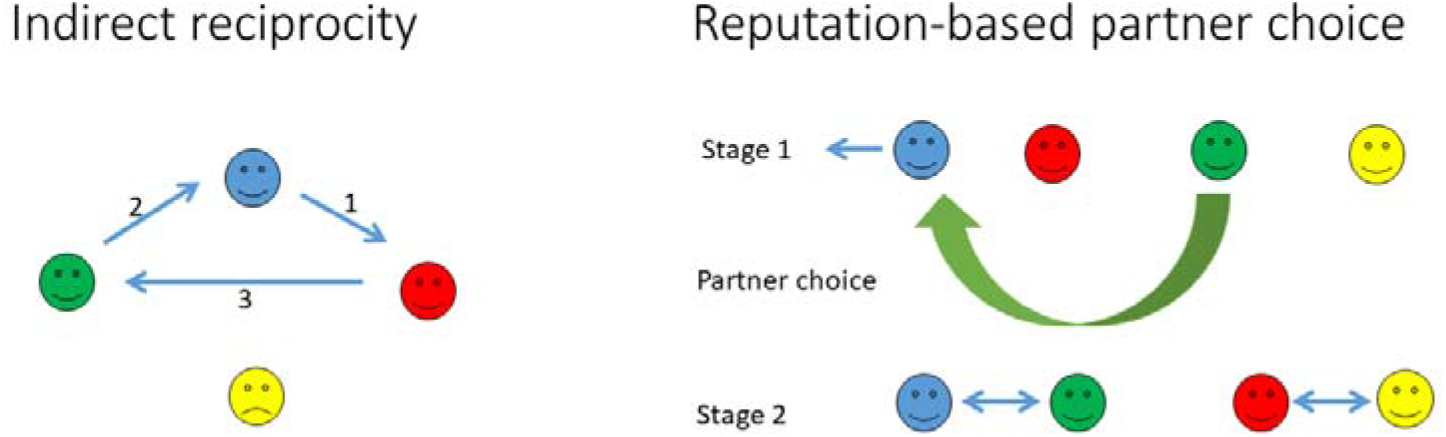
Mechanisms for benefitting from being seen to help others. **A.** Indirect reciprocity. Cooperation in indirect reciprocity involves a series of one-off donations in fluid populations to those who have been seen to help others. Here, the individual represented in blue helps red, each blue arrow representing paying a cost to benefit the recipient. Green observes that blue has helped, and in consequence, helps blue. Likewise red then helps green. Yellow does not help anyone and in consequence is not helped. **B.** Reputation based partner choice. In a social setting, individuals are aware of others’ reputations, which are gained by being seen to perform a helpful act to any individual. Through partner choice, cooperators assort for mutually beneficial long-term relationships. Here, blue (playing in the Competitor role) performs an act with a cost to itself and a benefit to any other party, here represented by a blue arrow away from the other individuals in order to avoid confusion with direct or indirect reciprocity. As a result, blue gains a positive reputation and so green (playing in the Selector role) chooses blue as a partner for a mutually beneficial relationship (two-way arrow). Red (playing in the Competitor role) does not perform a costly act but yellow (playing in the Selector role) accepts any partner for further interaction. The theory holds that those performing helpful acts are honestly signaling that they will make cooperative partners, and so that it makes sense for those looking for partners to choose on the basis of this signal.

Some key differences between indirect reciprocity and reputation-based partner choice may be summarized as follows:

1. The two theories aim to explain very different phenomena: indirect reciprocity aims to explain cooperation in large scale societies where individuals may not re-meet. In contrast reputation-based partner choice aims to explain how acts of apparent selflessness such as donations to charity, heroism, courtship feeding form part of long term social and sexual partnerships with repeated interactions.
2. Reputation-based partner choice is explicitly about signaling whereas indirect reciprocity is not. A signal is an adaptive behavior which benefits the signaler by changing a receiver’s behavior. This means that the cost invested in the signal may or may not benefit one or more recipients of any benefit and that the receivers of the signal may or may not be the recipients of such benefits. In contrast, indirect reciprocity involves a donation of a benefit to a recipient at a cost to the actor.
3. Indirect reciprocity is explicitly conditional in that it can only operate where individuals help those who help others. Reputation-based partner choice has no such conditionality. Individuals help others as a signal of cooperativeness. They benefit from then being more likely to be chosen for profitable partnerships.
4. Reputation-based partner choice involves partner choice for profitable relationships. The basic theory of indirect reciprocity involves no partner choice. It can be extended only in as much as individuals could theoretically choose who to give to. It cannot be extended to involve choosing as a partner someone that one has given to since this would involve direct reciprocity, which is a different theory of how helping Is beneficial.

The two theories have been compared experimentally and reputation-based partner choice has been found to be more effective in inducing reputational concerns (Sylwester and Roberts 2013).

The theory of reputation-based partner choice explicitly incorporates Zahavi’s Handicap Principle (Grafen 1990, Zahavi 1997), a form of costly signalling theory (Gintis, Smith, and Bowles 2001) whereby the signal (cooperative reputation) reveals any underlying differences in some quality such as resources. Inherent in this approach is that reputations are more than simply histories of how individuals have previously acted, each decision maximizing short term gains. Rather, reputations are costly investments and the challenge is then to determine how such investments can be adaptive. Standard costly signalling models applied to helping behaviour are based upon the notion that reputations allow observers to correctly deduce the underlying quality of the signaller (Gintis et al. 2001). An alternative or additional hypothesis is that costly investments might be honest signals of *intent*. That is, those who develop a reputation for helping might be more likely to be more cooperative and so may be preferred as partners (Roberts 1998; Van Vugt, Roberts, and Hardy 2007; Barclay 2013; Barclay 2015; Silk 2002; Bliege Bird and Smith 2005). However, the notion that reputations might signal future intent (rather than underlying quality) has been stated verbally by several of these authors yet has surprisingly not been developed theoretically. Thus, we do not have a coherent theory of when we can expect individuals to invest in reputations.

I consider investment in reputation to be a form of signalling (Maynard Smith and Harper 2003). The term ‘reputation’ is used to describe how other individuals judge someone based on their past behaviour. Reputation is explicitly linked to past helping behaviour in the model because reputation is formed when an individual indulges in helping. Helping behaviour in the model does not benefit an identified recipient either directly or indirectly so as to avoid any possible confusion with models of direct or indirect reciprocity. Reputation in the model is gained through a specific costly act and is not simply a record of past behaviour that may have reflected economic choices. In other words, we consider an act that is performed not because it brings reciprocal benefits, but because it changes the behaviour of other individuals. This means that reputation fulfils the criteria of being a signal and not just a cue (Maynard Smith and Harper 2003).

Here, I develop a model of how individuals choose partners for cooperative relationships. The aim is to show when it can pay to choose a social (or sexual) partner based on costly help, and consequently when helping provides a signal of intent to cooperate (trustworthiness). In the model, individuals compete for access to partnerships. They may or may not invest in a reputation by signaling, and signallers may or may not be more likely to be chosen as partners. Once partnerships are formed, the partners play rounds of a classical reciprocity game (Axelrod and Hamilton, 1981). In social dilemmas of this sort, defection can provide a short-term advantage, yet mutual cooperation pays when there are sufficient rounds within a partnership. It can therefore pay to select cooperative partners. The model aims to predict when signalling by helping others can be an honest signal of being a cooperative partner, i.e., when it pays to signal one’s cooperative intentions, trustworthiness, or commitment to a cooperative relationship, and when it pays to choose partners based on such signals.

## Methods

I used evolutionary computational simulation complemented by analytical work. The structure of the simulation models is outlined in Figure 2, while Figure S1 provides a more detailed flow diagram of the processes. Table S1 provides a reference list of parameters. The simulation was implemented in C++ in Microsoft Visual Studio 2017. Full programming code is provided in Supplementary Information.

**Figure 2.**
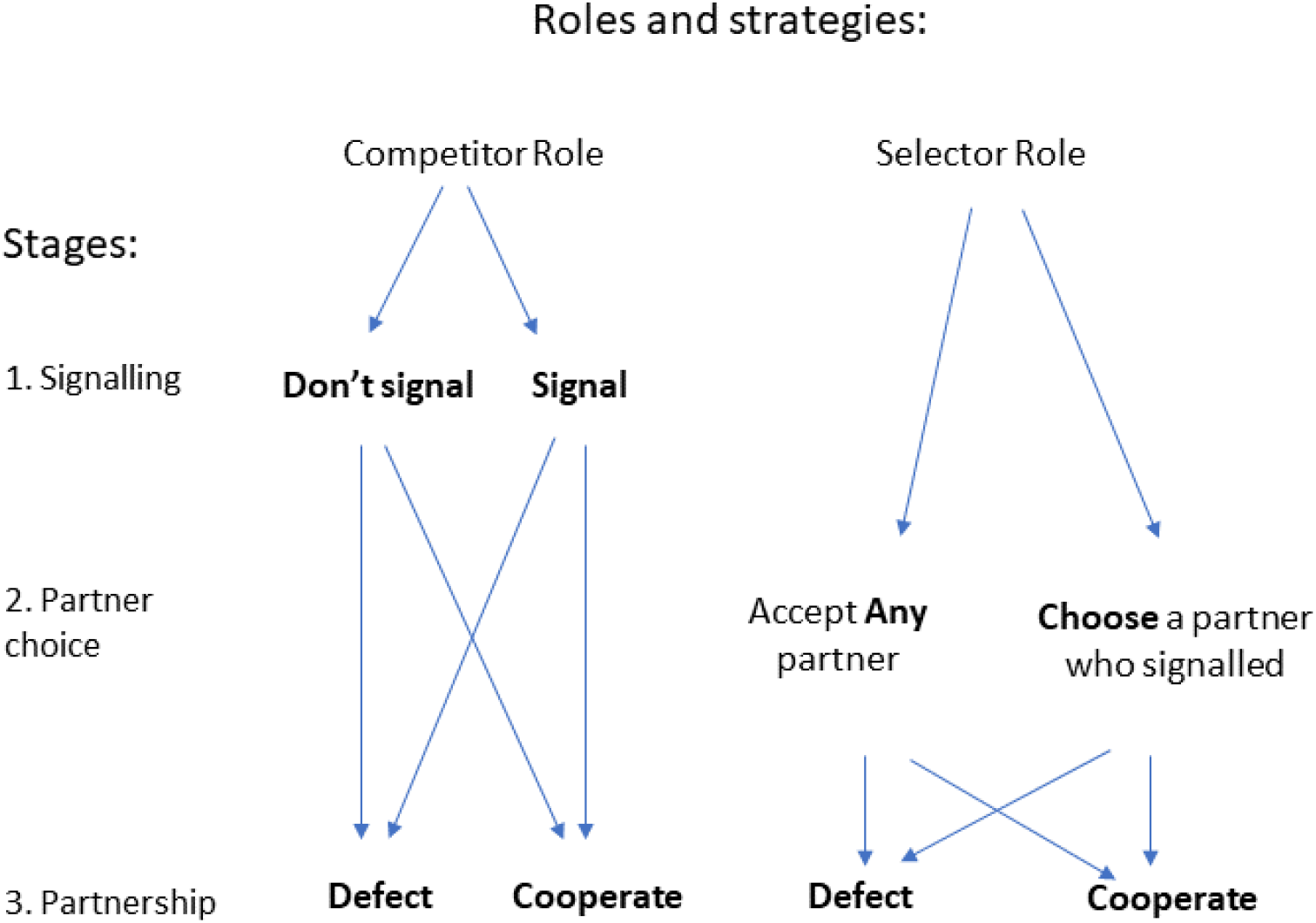
Roles and strategies. The model was based on the concept that some individuals designated as Selectors were seeking potential partners from among those designated as Competitors to form cooperative partnerships. The computer simulation model therefore began by dividing agents into those in the Competitor Role and those in the Selector Role. Competitors can either Signal, or they Don’t Signal (all strategy element names are capitalized and indicated in bold). Subsequently, Selectors can either accept Any partner generated at random by the program, or they can Choose a partner who signalled. Once paired, agents in either role may Cooperate or Defect. This structure specifies the eight strategies in the model, each being an independent combination of options represented by a pathway along the arrows shown e.g., a Selector may play a strategy of ‘Any’, ‘Cooperate’.

Agents were allocated by the computer program to one of two roles: those in the Competitor role competed to be selected to form partnerships with those in the Selector role. Selectors were assumed to be seeking partners to form a mutually beneficial relationship. Ahead of these partnerships, Competitors could pay a cost and thereby develop a ‘reputation’ in the eyes of a Selector, or they could opt not to pay this cost. Selectors could then decide whether to select those who had paid the cost (signalled) or to accept any partner presented to them by the program. Whenever Selectors accepted a partner, they then played a dyadic iterated social dilemma game in which reciprocity led to mutual benefit (see ‘Play’ below). Through evolutionary simulation, I investigated the fate of strategies of investing in a cooperative reputation by giving unconditionally versus withholding help; of selecting a partner with a good reputation or of accepting an assigned partner; and for both roles, of cooperating versus defecting in subsequent dyadic interactions. Figure 2 summarizes these strategies.

Dividing agents into the two roles allows us to study the evolution of strategies of signalling and choosing without knowing precisely how strategies are expressed in the real world. We could allow all individuals to express both strategies, but this would give us 16 strategies representing all possible combinations of signaling/ not signaling; choosing/not choosing; playing C/D after signaling; playing C/D after choosing. Findings from this more complex approach are consistent with those of the simpler model and so are not presented here. Furthermore, the division into two roles is consistent with other studies in the field including models of the costly signalling of quality (Gintis, Smith et al. 2001, Jordan, Hoffman et al. 2016) and the ‘biological market’ literature where individuals must first be classed as ‘buyers’ or ‘sellers’ (Noë and Hammerstein 1994) (Barclay 2013).

The model assumes that individuals differ in their profitability as cooperation partners solely in terms of their strategy. It therefore does not consider type differences due to resources or abilities that form the basis of typical costly signalling models (Gintis et al. 2001). I further assume that these expressed differences are detectable by potential partners as a signal or reputation index; and that individuals can choose partners for future interactions based on this. Crucially, there is no pre-set link between displaying and continuing to cooperate after being chosen and thus cheating by signalling and then exploiting a partner is possible in the system.

## Strategies

Agents in Selector and Competitor roles were assigned strategies by the program. Competitors were given strategies of either investing in a Signal or not (No Signal). Selectors had strategies of either choosing those who had invested in a Signal (Choose); or accepting all assigned partners (Any). Agents in both roles also had strategies of either cooperate (C); or defect (D) with the partner. Strategies were assigned independently, so that all four combinations of the two binary decision rules could evolve independently. The resulting roles and strategy combinations are summarized in Figure 2.

## Model Structure

The model employed a meta-population structure in which a default of *N* = 100 agents in each of the two roles was distributed on each of a default of *i* = 10 islands (hence a population size of 2000). Islands were units of reproduction (see below). Island structure was employed because it is thought to overcome problems with unrepresentative results arising in single populations (Leimar and Hammerstein 2001). Agents interacted with other agents within their islands.

## Initialization

At the start of each simulation, all Competitor agents were initialized with strategy elements set to “No Signal, Defect”, while Selector agents were initialized with “Any, Defect”, on the basis that we were interested in whether investing in signalling, and choosing signallers, can become established through association with cooperation in an environment which started with none of these. Agents in the Competitor role were initialized with a reputation index of neutral, reflecting the fact that each agent starts a generation with no interaction history. All agents started with a baseline payoff set to 1000 so that no payoffs would be negative and selection would be weak.

## Signalling and partner choice

The program simulated a process by which Selectors formed partnerships from among the Competitors. At the start of each generation all Competitor agents carrying the Signal strategy element paid the cost of signalling (i.e their baseline payoff was reduced by amount *s*) and their reputation index changed from ‘neutral’ to ‘positive’ (Figure 2, Stage 1). The program then cycled through all Selectors and assigned them partners at random from amongst the Competitors. If the Selector had the strategy ‘Any’ then a partnership was formed with the first randomly assigned Competitor. If the Selector had the strategy ‘Choose’ then it rejected randomly assigned partners until assigned one that did carry the Signal strategy element. Any Competitor that had previously been in partnership with a Selector was rejected. This meant that there was no opportunity for learning and memory from experience about which Competitors were cooperative, and therefore that partner choice could not involve reciprocity. All selections were subject to a giving up time of twice the population size (to avoid endless loops if no suitable partners could be found). This meant that should no suitable partner be found, the Selector had to settle for the last Competitor randomly assigned by the program. This process resulted in all Selectors having one interaction for each cycle *m* through the group, while Competitors had an average of one but had a variance depending on the extent of partner choice. Choice was costly in that Choosy agents faced a probability *h* that they would get no partner. This was implemented as a simple exogenous expected opportunity cost so that choosiness could not drift when signalling was common, but that further assumptions about how the costs of choice might vary were not made. By default, each island metapopulation was cycled through *m*=50 times, meaning that each agent had a 0.5 probability on average of meeting each other agent.

## Play

Once partnerships were formed, the agents played an iterated Prisoner’s Dilemma game. Agents carried strategies of ‘Cooperate’ (C) and ‘Defect’ (D). Cooperators played a simple conditional strategy of Cooperate until the partner defected (sometimes called “Grudger”). This meant that if both were C agents then they both cooperated on all rounds. Cooperation incurred a cost of *c*=1 payoff unit, with a benefit of *b*=2. However, if both agents were of type D, then no cooperation occurred in any rounds. If one agent played D and the other played C, then the former received a benefit whilst the other paid a cost. Agents that were exploited in this way responded by failing to cooperate in any further rounds (the Grudger strategy). The iterated game therefore had payoff matrix as follows in each round (payoffs are to the row player):

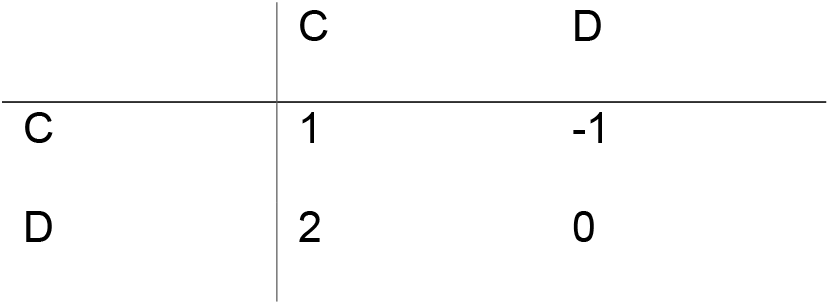

Given this payoff matrix, mutual cooperation could be profitable with repeated rounds and conditional strategies (like Grudger). Below a critical number of rounds *r*, Defectors would have the higher payoff, while above this value, Cooperators had a higher payoff. This is the classic game structure and result of (Axelrod and Hamilton 1981). It implements the basic conundrum of social dilemmas in that there can be a benefit in selecting cooperative partners for long term relationships, yet a risk that partners will defect and exploit cooperative first plays.

Agents had relationships with multiple other agents. In each of *m* meetings an agent followed the processes described above of competing for / selecting partners and then playing the repeated social dilemma with those partners (see program structure in Figure S1). As described in the subsection ‘Signalling and partner choice’, agents could not be paired with those with which they had already had a relationship. Therefore the model considered choice of partners for long term reciprocally beneficial relationships (within which memory must occur) yet was uncontaminated by memory for prior partners which could have led to reciprocity between relationships.

## Reproduction

Generations were discrete. At the end of each generation, strategy frequencies for the following generation were assigned in proportion to their success in the previous generation using the following method. The program cycled through all agents in each of the two roles separately. Payoffs were summed across all agents having each strategy in turn. This was done within each island and then across all islands, resulting in local and global strategy payoffs. Agents were assigned strategies in the next generation in proportion to the relative payoff of each strategy. An individual for the next generation was derived locally (i.e. within the island) with probability 0.8 and globally with probability 0.2. This followed the process recommended to reduce the potential for genetic drift and allow migration of successful strategies to other islands (Leimar and Hammerstein 2001). Population size was kept constant, with each discrete generation being fully replaced by the next. Reproduction was accompanied by mutation: with probability *μ*= 0.02 an individual’s strategy was replaced at random (any strategy could change to any other strategy without having to track through an imposed sequence). By default, simulations were run for 1000 generations and 10 replicates were run for each set of conditions. Simulations therefore allowed strategy frequencies to evolve across generations, with those strategies accumulating the highest payoff in one generation becoming more numerous in the next. All strategy combinations for both roles could evolve independently.

## Results

The aim of the simulation model was to determine whether a link between investment in a costly act, ‘reputation-building’, could become linked to cooperating, so that agents selecting partners would use reputation-building as a signal which provided information about future cooperative behaviour.

Considering first the evolutionary dynamics of all strategies, agents in the Competitor role evolved to invest in the signal and then cooperate (Figure 3a), while those in the Selector role evolved a tendency to choose Signallers preferentially and also to cooperate with their chosen partner (Figure 3b). Summing these strategies shows that signalling, choosing and cooperating all evolved readily from an environment of no signalling, accepting any partner and defecting (Figure 3c).

**Figure 3.**
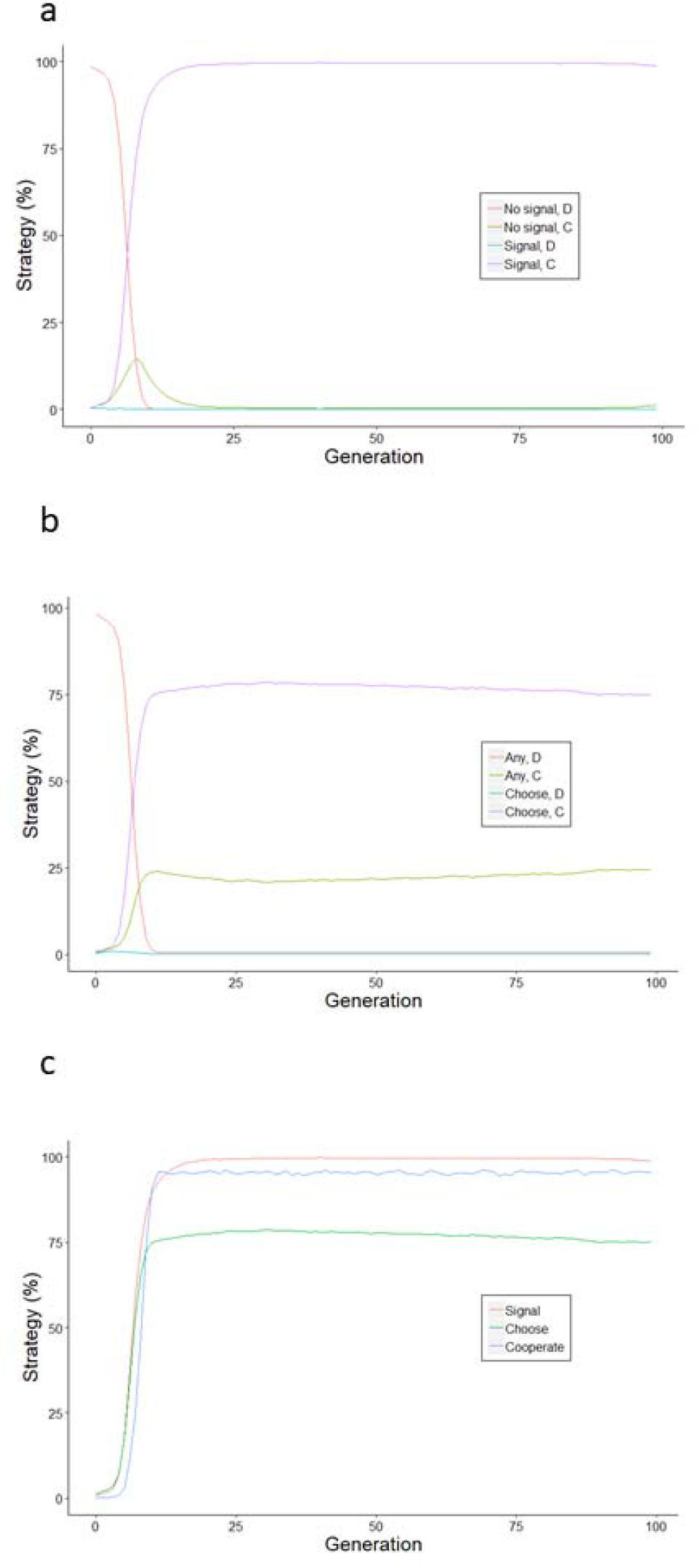
Reputation building evolves. Strategy dynamics are plotted for the first 100 generations (as strategy frequencies remained stable during the remaining generations of the 1000 for which the simulations were run) and are averaged across 10 simulations. (a) ‘Competitors’, showing all strategy combinations of ‘No Signal’, ‘Signal’ in Stage 1, and ‘C’ (Cooperate) or ‘D’ (Defect) in an iterated simultaneous Prisoner’s Dilemma in Stage 2. (b) ‘Selectors’, showing all strategy combinations of accept ‘Any’ or ‘Choose’ an agent that has signalled and whether they go on to cooperate with partner (as for part a). Parameters were population size *N* = 100; generations = 1000; islands *i* = 10; c=1; b = 10; meetings *m* = 50; rounds *r* = 25; signal cost *s* = 10; cost of choice *h* = 0.01; baseline payoff = 1000; mutation rate *μ*=0.02 (see Table S1).

Cooperators were more likely to signal than were defectors: the proportions of cooperators and defectors that signalled were 0.96 ± 0.01 and 0.42 ± 0.02 respectively (averaged over 10 simulations, of 100 generations; Figure 4a). Thus, those agents which went on to cooperate in stage 2 were much more likely to have signalled in stage 1. Looking at this from the perspective of a Selector, the probability that a Signaller was a Cooperator was much higher than the probability that a Non-signaller was a Cooperator (0.99 ± 0.00 and 0.65 ± 0.02 respectively; values from 10 simulations and 100 generations; Figure 4b)). This meant that choosing a Signaller rather than a Non-Signaller increased the chance of a Cooperative partner by approximately 52%. This probability was much greater when Signalling and Cooperation were becoming established whereas the quoted statistics are heavily weighted by the equilibrium conditions. These probabilities show that a link evolved between Signalling and Cooperation so that it paid to choose Signallers because they were more likely to be Cooperative partners. Reputation-based partner choice therefore worked when an association evolved between signalling and going on to cooperate with a partner, and therefore Selectors did use the signal in partner choice. It appears that when Cooperators are better able to engage in a costly act of helping others than are Defectors, this ability to engage in a costly act becomes useful information for Selectors, and hence in turn can promote engaging in costly acts by Cooperators in order to be chosen.

**Figure 4.**
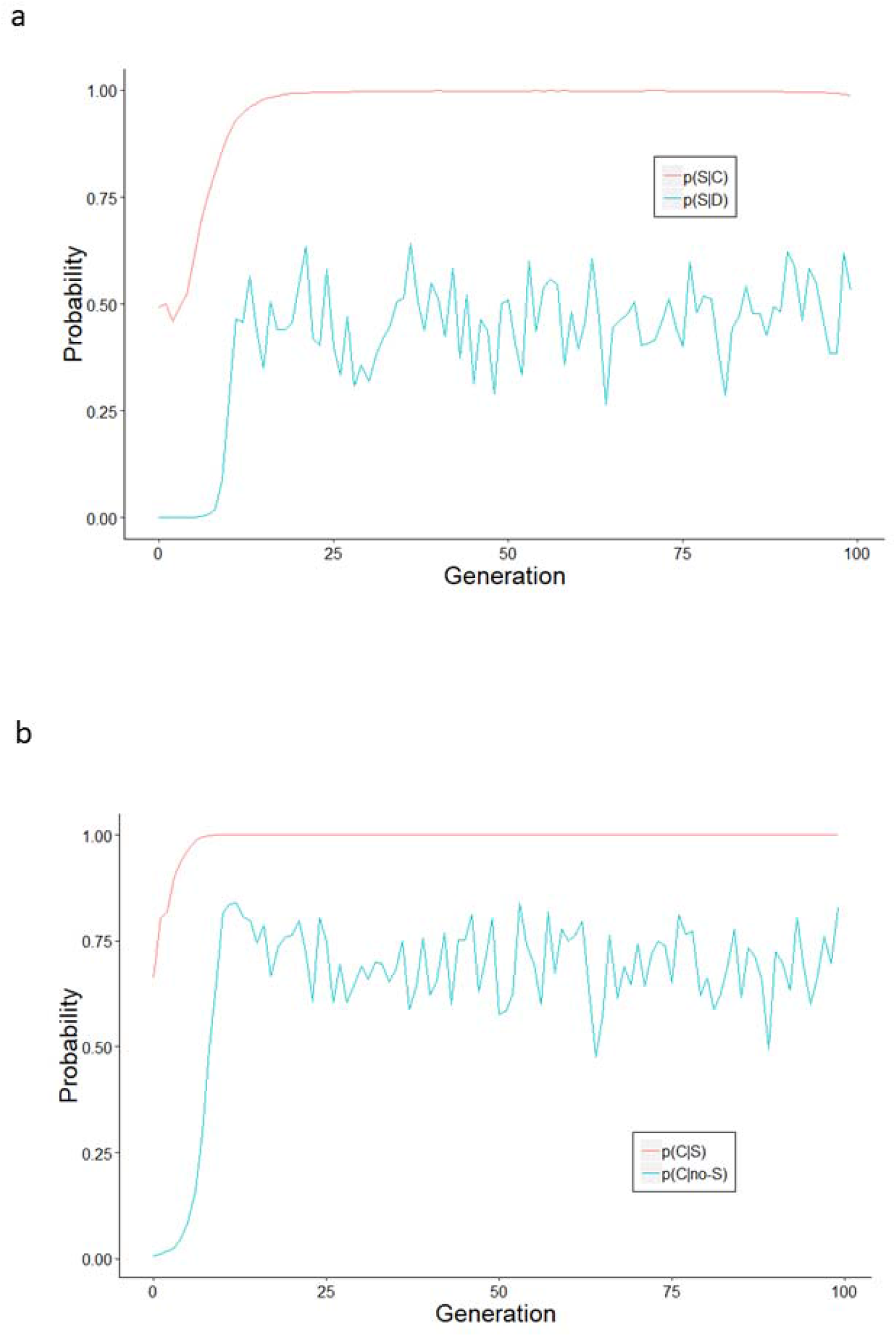
Evolutionary dynamics of signalling by cooperators and defectors. A. The probability of signalling given that an agent is a co-operator [p(S|C] versus a defector [p(S|D)]. B. The probability of being a cooperator given the agent is a signaller [p(C|S)] versus a non-signaller [p(pC|no-S)]. Simulation parameters as in Fig 3.

## Analysis

Why does investing in a good reputation (signalling) mean agents are chosen as partners, and how is the signal an honest indicator of future cooperation? To answer this we need to show how signalling, choosing and cooperating become linked. This will happen when the signal provides reliable information about the likelihood of an individual going on to cooperate. This means we need to find where it is strategic for cooperators but not defectors to signal. For signalling to be strategic, the costs of signalling *s* must be offset by the benefits of increased payoff when chosen. Simplifying by considering the payoff to cooperators and defectors on any one interaction when a signaller (who signals once) is chosen by a Selector who cooperates andwhen non-signallers achieve no interactions, then signalling should be strategic for cooperators when:

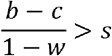

Where *b* and *c* are the costs and benefits of cooperation respectively and *w* is the chance of interactions continuing in an iterated prisoner’s dilemma game. And for defectors signalling is strategic when

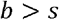

Thus, given these simplifications, signalling is strategic for cooperators but not defectors when:

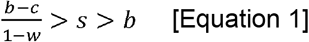

We can see from this result that there is a basic asymmetry between cooperation and defection strategies in the payoffs they get from being chosen; and hence that signalling provides information that the Competitor will go on to play Cooperate. Specifically, given these assumptions, signalling and cooperating will be linked when the payoff for signallers of maintaining reputation (in repeated rounds of mutual cooperation) is greater than the short-term payoff from exploiting good reputation. Conversely, it is not strategic for defectors to invest in a reputation (pay costs of signalling cooperativeness) and then exploit on the first round because they cannot recoup their signal costs when individuals remember they have defected within a repeated game.

This can be understood from Figure 6. Part (a) presents the typical costly signalling approach. This assumes that individuals differ in quality and this leads to differences in their stable levels of investment in signalling. A costly signal is then an honest indicator of underlying quality. This is the typical explanation for generosity: essentially, individuals that are very generous must be of high quality (e.g. Gintis et al 2003). Figure 5 (b) shows schematically the alternative theory presented here. My theory is that individuals differ in the benefits they get from each signal: provided Cooperators play a responsive strategy, they can establish partnerships in which they gain higher benefits from repeated rounds of cooperation than Defectors can gain by exploiting a Cooperator once. This can be interpreted as corresponding to the case in (Johnstone 1997a) where honesty in signalling is maintained due to differential benefits from signalling. However, instead of individuals differing in ‘need’ (as in chicks begging for food from their parents), individuals here differ in the benefits they can get by cooperating after signalling. Essentially, paying a high cost up front is an honest signal of going on to cooperate: individuals simply cannot afford to pay a high cost up front if they are simply going to go on to exploit responsive Cooperators once.

**Figure 5.**
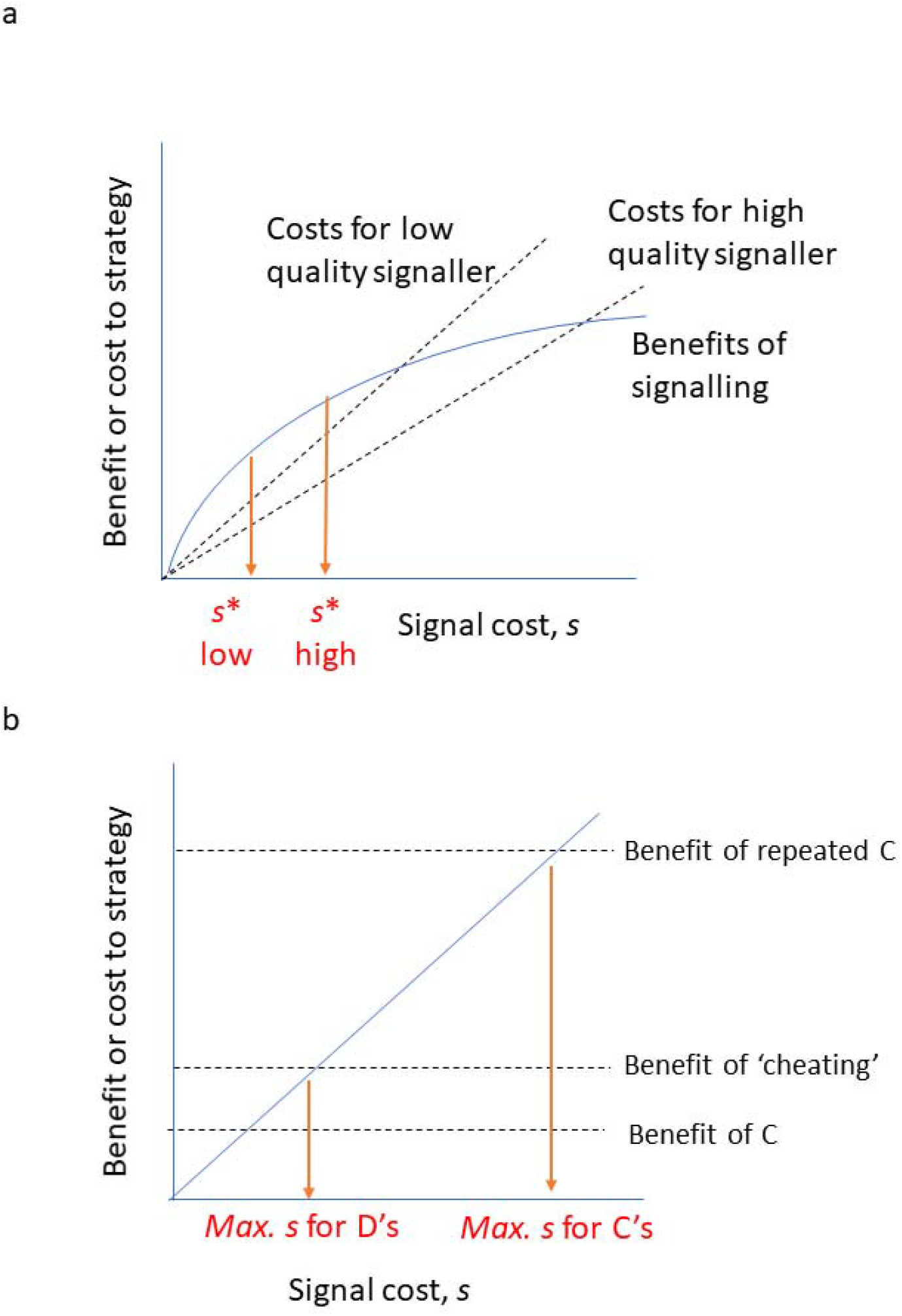
Signal honesty. The costs of signalling can maintain honesty in two ways. In (a), individuals belong to high and low quality classes. These classes differ in their investment in signalling and so the form of the signal indicates underlying quality. In (b) the costs of signalling are the same for all, but Cooperators get greater benefits from signalling when they engage in repeated interactions than do Defectors who can only exploit or cheat a partner once. The figure illustrates schematically how individuals playing repeated cooperation (C) can make a net profit despite paying signal costs, when this may not be possible for those ‘cheating’ i.e. defecting on co-operators. Between the maximum signal cost sustainable by D’s and the maximum sustainable by C’s is the region in which any Signaller must be a Cooperator. Adapted from (Johnstone 1997a) Fig 7.2a.

**Figure 6.**
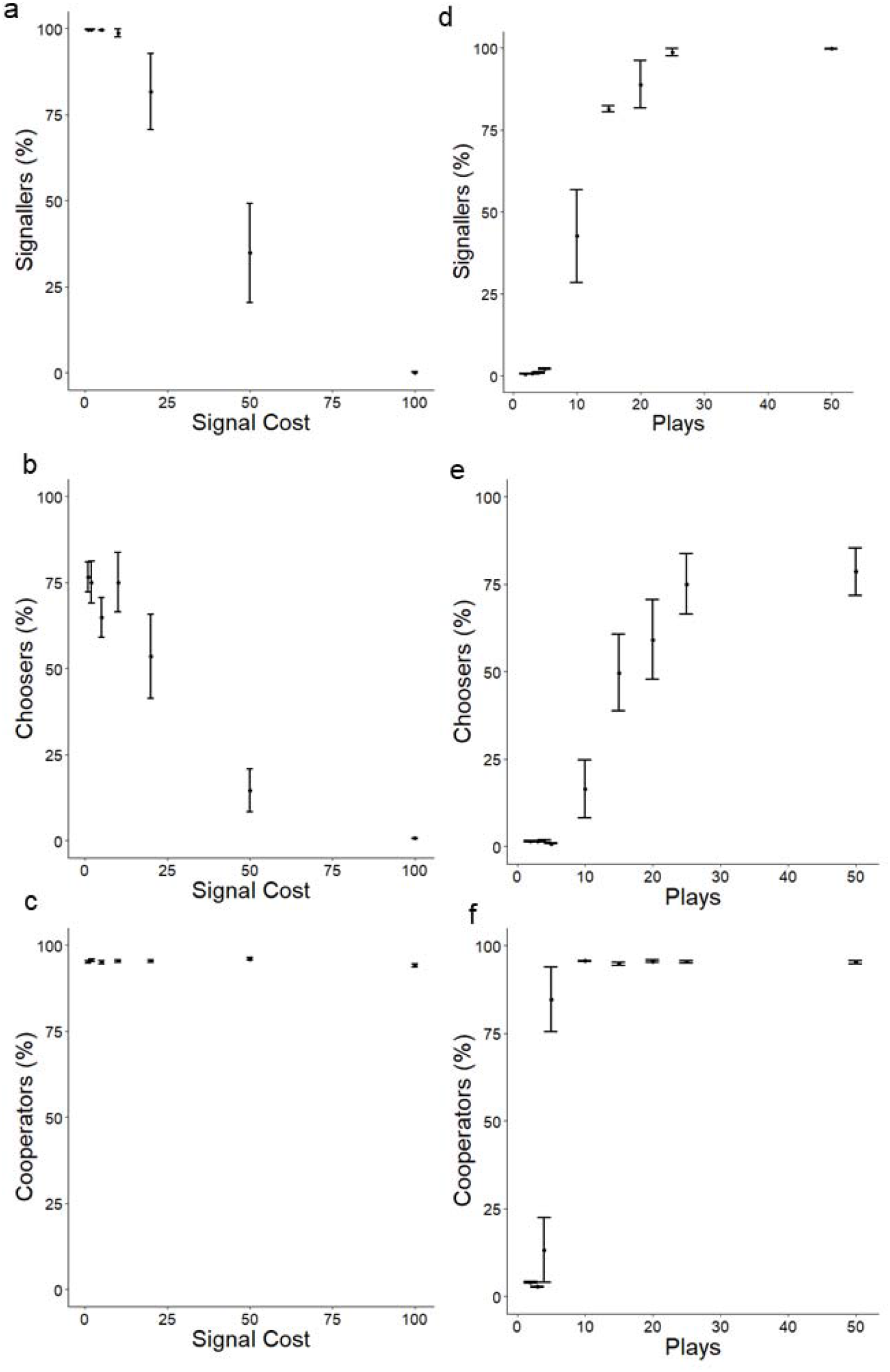
Effects of varying key parameters. Panels a-c show the effect of varying signal cost whilst keeping other parameters constant (see Figure 3 Legend for parameters) on the percentage of agents displaying choosy strategies; the percentage signalling; and the percentage cooperating. In each case these are calculated as means with standard errors of the given variable value in generation 100, computed across 10 simulations. They are therefore intended to indicate the average value at equilibrium (see Figure 3 for how the evolutionary dynamics reach equilibrium value within 100 generations). Similarly panels d-f show average percentages of Signallers, Choosers and Cooperators keeping other parameters constant (as in Figure 3) while varying the number of plays as the primary means of increasing the benefits of a relationship with a partner.

## Parameters

In the context of the simulations, the analysis predicts that Cooperators will have higher marginal return s investing in signalling than Defectors. The analysis further suggests there are two key factors affecting whether reputation-based partner choice will be found: on the one hand, the cost of the signal, and on the other the benefits of a cooperative relationship. Figure 6 displays the effects of varying the cost of signalling (a-c) and the number of plays within a partnership (d-f). At low signal costs, almost all agents evolve to invest in signalling and the majority use the signal in choosing a partner, but as signal cost increases, so the proportions of signallers and choosers declines. The proportion cooperating does not depend on signal cost (while the other parameters were kept constant). These results are qualitatively in line with the analytical prediction that signalling and choice will be found when signalling is honest, which will be when a low cost of signalling and a high benefit from repeated rounds make the marginal costs of signalling lower for co-operators than for defectors. For example, when the number of plays is high, there are larger gains to be made from cooperative partnerships and so co-operators have more to gain, hence it can be worth signalling that one is a co-operator.

I went on to investigate systematically the effects of departures from all default parameters used in the previously presented results by varying one parameter at a time and keep others constant. Increasing the cost of choice by increasing the proportion of times that choosy individuals missed out on partnerships led to a gradual reduction in both signalling and choice itself, but had little impact on cooperation (Figure S2). Increasing the benefit:cost ratio increased the long term benefits of cooperation and so as predicted increased the levels of signalling and choice (Figure S3). Increasing the population size, N, above the default level had little effect on signalling but choice fell away, probably because choice is proportionally less effective when searching through larger populations (Figure S4). Signalling and choice fall away at high numbers of meetings (Figure S5). This is because agents are allowed to meet any other agent only once, so partner choice becomes impossible as the number of meetings rises towards the total number of other agents to meet. Mutation rate had little quantitative impact on signalling, choice or cooperation, showing that the conclusions are not dependent upon a particular level of variation (Figure S6). The baseline value for payoff used did not impact on behaviour, showing the results are robust to the strength of selection (Figure S7). There was some variability in results at low numbers of islands, but at or above the default value of 10, there was little further quantitative impact of varying population structure in this way (Figure S8).

## Discussion

These results illustrate how paying a cost to help others can be strategic for those who go on to form mutually beneficial cooperative interactions and not for those who take the short-term benefit of defection. They show that the differential benefits of cooperating and defecting strategies can lead to costly helping behaviour being an honest signal of future commitment to cooperation (trustworthiness). Reputation-building provides information about strategy, based simply on the fundamental difference between cooperators and defectors. Reputations evolve into honest signals of commitment which it pays to attend to when seeking cooperative partners. The results of the model are therefore in line with prior experimental findings that costly investment in reputation-building can bring net benefits through access to profitable partnerships (Sylwester and Roberts 2010). The conditions in which this is found are given by Equation 1. In particular, there will be a trade-off between the cost of signalling and the benefits of iterated cooperation, such that higher signal costs can be supported when seeking to engage in long, profitable relationships. This makes important predictions about when we can expect to see reputational displays in humans and other animals, whether in courtship, in social interactions and in marketing.

The results can be interpreted as showing how partner choice creates assortment among co-operators (e.g. (Sherratt and Roberts 2012, Wang, Suri et al. 2012, Nax and Rigos 2016). The success of cooperation depends upon the act of cooperating leading to an increase in the chance of receiving help. By honestly signaling that they will cooperate, agents in the model assort with other co-operators.

Reputation-based partner choice contrasts with existing attempts to explain cooperative displays. Considering the popular notion of indirect reciprocity (which depends on a conditional rule of “help those who help” on each play), the two-stage structure employed here as a solution to social dilemmas (Roberts 1998) means that the costs of reputation building in one game are offset by access to profitable partnerships in a second. This crucial difference allows for unconditional help (e.g. charitable giving, public goods contributions); it shows how helping is in an individual’s strategic interests (compared to indirect reciprocity where there is a conflict between increasing image score and demonstrating discrimination (Leimar and Hammerstein 2001); and it requires no complex ‘moral’ rules of who to give to, such as the ‘leading eight’ (Ohtsuki and Iwasa 2006).

The reputation-based partner choice model also differs from current applications of costly signalling theory to reputations. These models are based on finding a separating equilibrium between two types differing in underlying quality, whether genetics, health, resources or competitive ability (Gintis, Smith, and Bowles 2001). Costly signal honesty occurs when it signalling is condition dependent, so that low quality individuals suffer higher marginal costs or lower marginal benefits from signalling. In contrast, any individuals in the reputation-based partner choice model could play strategies of cooperator or defector (there are no quality differences) and the frequencies of these strategies evolve in proportion to payoffs (as opposed to simply being assumed to exist as separate types as in costly signalling models). In the reputation-based partner choice model, signalling then provides information about strategy itself (not quality type) because cooperative strategists have a comparative advantage (Ricardo 1891). In contrast to a recent model of third party punishment as a signal of trust (Jordan et al. 2016), the reputation-based partner choice model does not depend on unsubstantiated assumptions. In particular, there is no assumption that individuals belong to fixed trustworthy or exploitative types that cannot change in frequency due to selection, nor that trustworthiness is associated with lower net costs of third-party punishment.

It is sometimes stated that a cooperative reputation indicates both ability and intent to cooperate (Van Vugt, Roberts, and Hardy 2007; Barclay and Willer 2007), yet costly signalling theory focuses on the former (Gintis, Smith, and Bowles 2001). In contrast, the model presented here is about signals of intent, or strategy, with no necessary difference in ability. Intention signalling is honest because those going on to a cooperative partnership can afford to invest in signalling whereas those that exploit co-operators will get a high payoff once, but this is likely to be insufficient to make signalling profitable. By signalling, agents demonstrated that they were investing in a long term relationship; and so it paid those choosing partners to prefer signallers. This model provides theoretical evidence in support of the hypothesis that I had previously presented only verbally (Roberts 1998). The concept of signalling intentions is being recognized by a few authors as being important in explaining human behaviour, for example in Martu hunters where “signals may be crucial for ensuring trust and commitment with long-term partners” (Bird et al. 2018), and more generally in hunter-gatherers where “although the focus of much traditional costly signaling literature has been hunting prowess [there is also] the potential of hunting to signal generosity and garner cooperation partners” (Stibbard Hawkes 2019).

The issue of maintaining reputation versus taking a short term temptation is analogous to the condition for reciprocity in an iterated prisoner’s dilemma (Axelrod and Hamilton 1981). When the ‘shadow of the future’ is large, then it pays to play Tit-for-Tat in repeated games, i.e. where (*b*-*c*)/(1-*w*) > *b*. Simply changing the benefit of defection (*b*) to be the cost of signalling results in the simplified equation derived above for when it pays to cooperate. Thus in reciprocity it pays to start by cooperating when there are sufficient benefits of repeated interaction, and analogously in reputation building it pays to start by signalling when this cost is exceeded by the benefits of being chosen for repeated cooperation.

As we have seen, the reputation-based partner choice model is based on paying an upfront cost. In this it is convergent with the notion of commitment devices (Frank 1988). The rationale is that an initial investment commits one to a course of action, in this case benefiting from long term cooperation as opposed to taking the temptation to defect. One can predict that the greater the potential benefits from cooperative relationships, the greater the up-front cost that can be sustained in signalling commitment [compare recent work on commitment in language (Vullioud et al., 2017). However, the reputation-based partner choice model also differs from a model of ‘cheap talk’ where low cost signals can indicate intent provided there are repeated interactions (Silk, Kaldor, and Boyd 2000) in that it specifically explains costly reputation-building and in that reputations are the result of single signalling events rather than being learned over many rounds.

The work presented here necessarily has limitations in its application. Each signalling system in the real world may have different properties. The model is intended to apply to behavioural signals. Individuals are envisaged to perform some behaviour directed towards observers who may or may not elect to respond to the behaviour. Specific models of real-world systems would need to tailor the model to account for factors such as how many potential partners can be influenced by one signal.

Furthermore, while this model follows standard practice in dividing individuals into two roles (here Competitors and Selectors while the Biological Market literature considers ‘buyers’ and ‘sellers’), further work should model the case where individuals can perform both roles.

In conclusion, I have shown that introducing signalling changes the dynamic between cooperative and defecting strategies because cooperators gain larger marginal benefits from investing in signalling to a potential partner than do defectors. Reputation itself provides an honest signal of long term commitment, or trustworthiness, providing a means by which cooperators can select each other for mutually beneficial relationships. The development of this theory is important in adding substance to previous verbal acknowledgements that signals might indicate future strategy in addition to giving information about quality or resources (Roberts 1998; Van Vugt, Roberts, and Hardy 2007; P. Barclay 2013; Pat Barclay 2015; Silk 2002; Bliege Bird and Smith 2005). The formal theory of honest signalling of cooperative intentions that I have advanced here fits well within recent verbal arguments for such a theory (Bliege Bird and Power 2015, Bird, Ready et al. 2018, Stibbard-Hawkes 2019)

Tests of the reputation-based partner choice model have already been carried out based on the verbal two-stage structure outlined in (G. Roberts 1998), and so the model seeks to offer a formal explanation for existing experimental results showing the signalling function of helping as a signal of trustworthiness (Sylwester and Roberts 2010; Barclay 2004; Fehrler and Przepiorka 2013) and commitment (Yamaguchi, Smith, and Ohtsubo 2015). Nevertheless, future experiments using economic games should test the quantitative predictions of the model while varying its parameters. The reputation-based partner choice model forces us to think carefully about whether signals indicate differences in underlying quality or in strategy (although of course in practice the two may intertwine). Individuals may reveal their long term cooperative strategy by helping, by donating to charity or contributing to public goods; brands may display long term strategy through advertising investment; primates may demonstrate investment in social bonds; monogamous birds such as albatrosses may give commitment signals where long relationships are decisive. The key factor is that reputation-building can signal that individuals are committed for the long term.

